# Novel circadian clock activators display anti-obesity efficacy via suppression of adipocyte development and hypertrophy

**DOI:** 10.64898/2026.01.08.698495

**Authors:** Xuekai Xiong, Jemima Pangemanan, Tali Kiperman, Lu Tang, Zhipeng Fang, Alon Agua, Wendong Huang, David Horne, Ke Ma

## Abstract

The circadian clock exerts temporal coordination of metabolic processes to maintain homeostasis, and its disruption predisposes to the development of obesity and insulin resistance. Despite the established genetic basis of clock modulation in adipocyte development, whether it can be targeted for anti-obesity interventions remains to be explored. Here we report the novel actions of clock-activating molecules, chlorhexidine and a new derivative CM002, on inhibiting adipocyte development and hypertrophy that results in anti-obesity efficacy *in vivo*. Both chlorhexidine and CM002 were sufficient to activate clock in adipocytes with induction of core clock components and shortening of clock period length. Consistent with their clock-activating properties, these compounds suppressed the distinct lineage commitment and terminal differentiation stages of adipogenic precursor cells mediated via activation of the Wnt signaling pathway. Furthermore, CM002 attenuated lipid storage and adipocyte hypertrophy by suppressing the lipogenic and adipogenic program in a clock-dependent manner. Most importantly, CM002 administration in mice with diet-induced obesity was sufficient to induce clock activation in adipose depots, leading to robust suppression of adipogenic factors and lipogenic enzymes with marked effect on reducing fat mass and promoting insulin sensitivity. Collectively, our findings uncovered the anti-adipogenic properties of novel small molecule clock activators with demonstrated anti-obesity efficacy. These compounds provide novel chemical probes to dissect clock function in metabolic regulations with translational potential toward development of first-in-class clock-targeting drugs for anti-obesity therapy.

**Highlights:**

- Discovery of the anti-adipogenic properties of the clock activator chlorhexidine
- Identification of a new clock-activating molecule CM002
- CM002 inhibits the lineage commitment and terminal differentiation of adipocytes
- Clock activation by CM002 suppresses lipid storage in mature adipocytes
- CM002 displays anti-obesity efficacy in diet-induced obesity model

## 1. INTRODUCTION

Circadian clock, a molecular machinery that drives ∼24 hour oscillations, exerts pervasive temporal regulation in diverse metabolic processes [1, 2]. Through orchestration of time-of-day dependence of key metabolic pathways in distinct organs, circadian control is required to maintain metabolic homeostasis [2]. Disruption of this timing mechanism, increasingly prevalent in our modern lifestyle, leads to metabolic dysregulations contributing to the development of obesity and Type II diabetes [3–5]. Accumulating evidence from large-scale epidemiological observations and experimental investigations to date have established the intimate link between circadian misalignment and metabolic disorders [4, 6–10].

The cell-autonomous circadian clock is generated by a transcriptional/translational feedback circuit [1] composed of positive and negative regulatory components. Transcription activators CLOCK (Circadian Locomotor Output Cycles Kaput) and Bmal1 (Brain and Arnt-Like 1) initiates clock transcription that is countered by the feedback inhibition of their direct target genes and transcription repressors, Periods (Per 1-3) and Cryptochromes (Cry1 & 2). In addition, RORα (Retinoid-related Orphan Receptor α) and Rev-erbα, a pair of antagonistic transcription factors, exerts positive and negative regulations respectively, to control Bmal1 transcription that forms an re-enforcing loop of the clock mechanism [11, 12]. Genetic models of both the positive and negative arms of the core clock feedback loop indicate that the clock circuit is involved in modulating adipogenic processes. *CLOCK* mutant or ablation of *Bmal1* in mice resulted in the development of obesity [13–15]. Bmal1 exerts direct transcriptional activation of key components of the Wnt signaling pathway in adipogenic progenitors that suppresses differentiation [13], while PER2 is known to repress PPARγ that inhibits adipocyte development [16]. In mature adipocytes, clock controls rhythmic lipolytic enzyme gene expression that modulates lipid mobilization and consequently accumulation of triglyceride and adipocyte hypertrophy [17]. Given the key roles of clock in regulating adipocyte biology, targeting clock function to suppress adipocyte development and maturation may provide novel avenues to treat obesity and related metabolic consequences [18–20].

In an obese state such as high-fat diet-induced obesity, significant dampening of clock oscillation amplitude was evident [21], while nutritional overload and clock disruption were found to be synergistic in inducing diabetes [22]. On the other hand, time-restricted feeding could counteract high-fat diet-induced clock dampening that yields metabolic benefits [23]. Thus, maintaining or enhancing the metabolic outputs of circadian clock could be pursued for metabolic disease therapy. Notably, activation of the clock repressor Rev-erbα and a direct CLOCK/Bmal1 target by specific agonists, SR9009 and SR9011, demonstrated their anti-obesity effects in mice with improvement of dyslipidemia [24]. A recent study revealed that RORα modulation by a natural ligand Nobiletin enhanced the oscillatory amplitude of circadian clock [25]. We found that this RORα-activating flavonoid compound demonstrated robust inhibition of adipogenesis and adipocyte hypertrophy, resulting in anti-obesity effect in vivo with marked loss of fat mass [26]. However, despite the increasing effort to identify clock-modulatory small molecules for drug development [25, 27–29], targeting clock function in adipocytes for anti-obesity applications remains unexplored.

Based on the circadian clock regulation of key processes involved in adipocyte maturation and lipid storage, here we show that augmenting its function by novel activators, via a recently identified molecule chlorhexidine (CHX) or a new structural analog CM002, resulted in strong inhibition of distinct stages of adipocyte development [30]. Most importantly, in vivo studies of CM002 revealed its robust anti-obesity efficacy, demonstrating the potential for developing clock-activating molecules for metabolic disease interventions.

## 2. METERIALS & METHODS

### 2.1 Animal studies

Mice were maintained in the City of Hope vivarium under a constant 12:12 light dark cycle provided with nestlets and maze for cage enrichment. All animal experiments were approved by the Institutional Animal Care & Use Committee (IACUC) of City of Hope and performed according to the IACUC approval. C57BL/6J mice were purchased from Jackson Laboratory and used for experiments following 2 weeks or longer of acclimation at 12 or 20 weeks of age. The mice on chow diet at 20 weeks of age were given an intraperitoneal injection for three doses per week up to 18 days with CM002 (5 mg/kg) or vehicle (DMSO). For high-fat diet feeding, mice at 14-20 weeks of age were fed with 45% high-fat diet (D12451, Research Diets) for 8 weeks, before daily IP injection with CM002 (5 mg/kg) or vehicle for 7 days, or CM002 (7 mg/kg) or vehicle three doses per week for 16 days. Adipocyte-specific Bmal1-null mice were described previously [31]. Mice were maintained in the lab and used for primary preadipocyte isolation at 10-12 weeks of age. For hematoxylin and eosin histology, adipose tissues were dissected at the end of the fixed in 10% neutral-buffered formalin for 72 hours prior to embedding. 10 μm paraffin sections were processed for hematoxylin and eosin staining.

### 2.2 Cell culture and adipogenic differentiation

3T3-L1 and C3H10T1/2 cell lines were obtained from ATCC and maintained in DMEM with 10% fetal bovine serum supplemented with 1% Penicillin-Streptomycin-Glutamine, as previously described [32, 33]. 0.25% Trypsin was used for digestion and subculture of these cell lines. For adipogenic differentiation, induction media containing 1.6μM insulin, 1μM dexamethasone, 0.5mM IBMX and 0.5 uM Rosiglitazone was used for 3 days followed by maintenance medium with insulin for 3 days for 3T3-L1, and for 5 days for C3H10T1/2 cells, as previously described [26, 33].

### 2.3 Generation of stable adipogenic progenitor cell lines containing Per2::dLuc

3T3-L1 preadipocytes and C3H10T1/2 mesenchymal stem cells obtained from ATCC were used for *Per2::dLuc* lentiviral transduction and stable clone selection using puromycin, as described previously [13, 26]. Briefly, cells were transfected with lentiviral packaging plasmids (pSPAX.2 and pMD2.G) and lentivirus vectors *Per2::dLuc* using PEI Max (Polysciences). At 48 hour post-transfection, lentiviruses were collected. 3T3-L1 and C3H10T1/2 cells were infected using collected lentiviral media supplemented with polybrene. 24 hours following lentiviral infection, stable cell lines were selected in the presence of 2 μg/ml puromycin.

### 2.4 Primary preadipocyte isolation and adipogenic induction

The stromal vascular fraction containing preadipocytes were isolated from subcutaneous fat pads, as previously described [34]. Briefly, fat pads were cut into small pieces and digested using 0.1% collagenase Type 1 with 0.8% BSA at 37^0^C with constant shaking for 60 minutes. The digested homogenate was passed through Nylon mesh and centrifuged to collect the pellet containing the stromal vascular fraction with preadipocytes. Pelleted preadipocytes were cultured in F12/DMEM supplemented with bFGF (2.5 ng/ml), expanded for two passages and subjected to differentiation in 6-well plates at 90% confluency. Adipogenic differentiation was induced for 2 days in medium containing 10% FBS, 1.6 μM insulin, 1 μM dexamethasone, 0.5 mM IBMX, 0.5 uM rosiglitazone before switching to maintenance medium for 4 days with insulin only. Compounds at indicated concentrations were administered for the entire differentiation time course at day 4 following adipogenic induction.

### 2.5 Human primary preadipocyte isolation and adipogenic induction

Human primary preadipocytes were isolated from visceral fat of human donors obtained via the Arthur Riggs-Diabetes and Metabolism Research Institute Pancreatic Islet Cell & Tissue Processing Service Center at City of Hope. A similar protocol was used as for mouse preadipocyte isolation. Briefly, fat pads were digested, and the stromal vascular fraction was collected and cultured in F12/DMEM supplemented with bFGF (2.5 ng/ml). Cells were expanded for two passages and subjected to differentiation in 6-well plates at 90% confluency. Adipogenic differentiation was induced for 4 days in medium containing 10% FBS, 1.6 μM insulin, 1 μM dexamethasone, 0.5 mM IBMX, 0.5 uM rosiglitazone before switching to maintenance medium for up to 9 days with insulin and rosiglitazone. Compounds at indicated concentrations were administered for the entire differentiation time course following adipogenic induction or at later stage of 6 days after differentiation.

### 2.6 Oil-red-O and Bodipy staining

Staining for neutral lipids during adipogenic differentiation were performed as previously described [32]. Briefly, for oil-red-O staining, cells were fixed with 10% formalin and incubated in 0.5% oil-red-O solution for 1 hour. Bodipy 493/503 was used at 1mg/L together with DAPI for 15 minutes, following 4% paraformaldehyde fixation and permeabilization with 0.2% triton-X100.

### 2.7 Continuous Bioluminescence monitoring of Per2::dLuc luciferase reporter

3T3-L1, C3H10T1/2 or primary preadipocytes containing a *Per2::dLuc* luciferase reporter were used for bioluminescence recording, as previously described [26]. Cells were seeded at 4×10^5^ density on 24 well plates at 90% confluence following overnight culture with explant medium luciferase recording media. Explant medium contains DMEM buffer stock, 10% FBS, 1% PSG, pH7 1M HEPES, 7.5% Sodium Bicarbonate, Sodium Hydroxide (100 mM) and XenoLight D-Luciferin bioluminescent substrate (100 mM). Raw and subtracted results of real-time bioluminescence recording data for 6 days were exported, and data was calculated as luminescence counts per second, as previously described [35]. LumiCycle Analysis Program (Actimetrics) was used to determine clock oscillation period, length amplitude and phase. Briefly, raw data following the first cycle from day 2 to day 5 were fitted to a linear baseline, and the baseline-subtracted data (polynomial number = 1) were fitted to a sine wave, from which period length and goodness of fit and damping constant were determined. For samples that showed persistent rhythms, goodness-of-fit of >80% was usually achieved.

### 2.8 TOPFlash luciferase reporter assay

M50 Super 8xTOPFlash luciferase reporter containing Wnt-responsive TCF bindings sites was a gift from Randall Moon [36] provided by Addgene (Addgene plasmid # 12456). For transient transfection with luciferase reporter construct, cells were seeded and grown overnight to reach 90% confluency. 24 hours following transfection, 10% Wnt3a conditioned media obtained from L Wnt-3A cell line (ATCC CRL-2647) was added to induce Wnt signaling. Luciferase activity was assayed using Dual-Luciferase Reporter Assay Kit (Promega) in 96-well black plates. TOPFlash luciferase reporter luminescence was measured on microplate reader (TECAN infinite M200pro) and normalized to control FOPFlash activity, as previously described [13]. The mean and standard deviation values were calculated for each well using four replicates and graphed.

### 2.9 Immunoblot analysis

Total protein was extracted using lysis buffer containing 3% NaCl, 5% Tris-HCl, 10% Glycerol, 0.5% Triton X-10 in MilliQ water with protease inhibitor cocktail. 20-40 µg of total protein was resolved on 10% SDS-PAGE gels followed by immunoblotting on PVDF membranes (Bio-rad). Membranes were developed by chemiluminescence (SuperSignal West Pico, Pierce Biotechnology) and signals were obtained via a chemiluminescence imager (Amersham Imager 680, GE Biosciences). Primary antibodies used are listed in Supplemental Table 1.

### 2.10 RNA extraction and RT-qPCR analysis

PureLink RNA Mini Kit (Invitrogen) was used to isolate total RNA from cells. cDNA was generated using Revert Aid RT kit (ThermoFisher) and quantitative PCR was performed using SYBR Green Master Mix (Thermo Fisher) in triplicates on ViiA 7 Real-Time PCR System (Applied Biosystems). Relative gene expression was calculated using the comparative Ct method with normalization to 36B4 as internal control. PCR primer sequences were listed in Supplemental Table 2.

### 2.11 Indirect calorimetry

Whole-body energy homeostasis was analyzed using the Promethion Core Metabolic System (Sable Systems) by the City of Hope Comprehensive Metabolic Phenotyping Core. Mice were single housed and acclimated in metabolic cages using the Comprehensive Laboratory Animal Monitoring System (Columbus Instruments), with ad libitum food and water with controlled lighting for 1 day prior to metabolic recording. Metabolic parameters, including oxygen consumption, respiratory exchange ratio, ambulatory activity, and food intake were recorded for 3 consecutive days, as described. Mice were given an intraperitoneal injection daily with CM002 (5 mg/kg) or vehicle (DMSO) for 4 days.

### 2.12 Plasma metabolite analysis and Insulin tolerance test

Plasma levels of glucose (Thermo Scientific), triglyceride (Teco Diagnostics) and free fatty acids (BioAssay Systems) were measured using 5-10 μl of plasma samples with respective commercial kits, according to manufacturer’s protocols. Insulin tolerance test (ITT) was performed as previously described [37]. Mice were fasted for four hours prior to intraperitoneal injection of 0.5 U/kg insulin and glucose levels were measured using a glucometer (AimStrip Plus).

### 2.13 Statistical analysis

Data were presented as mean ± SD. Each experiment was repeated at minimum three times to validate the result. The number of replicates were indicated for each experiment in figure legends. Two-tailed Student’s t-test or One-way ANOVA with post-hoc analysis for multiple comparisons were performed as appropriate as indicated using GraphPad PRISM. P<0.05 was considered statistically significant.

## 3. RESULTS

### 3.1 Clock-activating properties of chlorhexidine and a new derivative CM002 promote function of adipocyte-intrinsic circadian clock

The circadian clock transcription activator, Bmal1, inhibits adipocyte development through its transcriptional control of the Wnt signaling pathway [13, 38], and our recent study identified CHX as a novel clock-activating molecule by promoting CLOCK/Bmal1 transcriptional activity [30]. We generated stable cell lines containing the *Per2::dLuc* luciferase in mesenchymal precursor C3H10T1/2 (10T1/2) and 3T3-L1 preadipocytes to examine CHX effect on cell-intrinsic clock function in these adipogenic progenitor models. In 10T1/2 mesenchymal precursor cells, CHX treatment at 1 and 2 μM led to a dose-dependent shortening of clock period length (Fig. 1A & 1B). In addition, CHX at these concentrations were able to augment clock cycling amplitude (Fig. 1C), consistent with its function as a clock activator. The effects of CHX on reducing period length and increasing amplitude were also observed in lineage-committed 3T3-L1 preadipocytes at 1 μM (Fig. S1A-C). Furthermore, in primary preadipocytes isolated from the *Per2:dLuc* knock-in mice [39], CHX exhibited a similar effect on shortening period length (Fig. 1D), although it did not alter clock amplitude (Fig. 1E). Analysis of clock gene regulation by CHX in 3T3-L1 preadipocytes revealed induction of *Bmal1* together with CLOCK/Bmal1 direct target genes, *Nr1d1*, in line with activation of CLOCK/Bmal1-mediated transcription (Fig. 1F). Largely similar results of CM002 up-regulation of core clock genes were also evident in 10T1/2 mesenchymal precursor cells (Fig. S1D).

**Figure 1.**
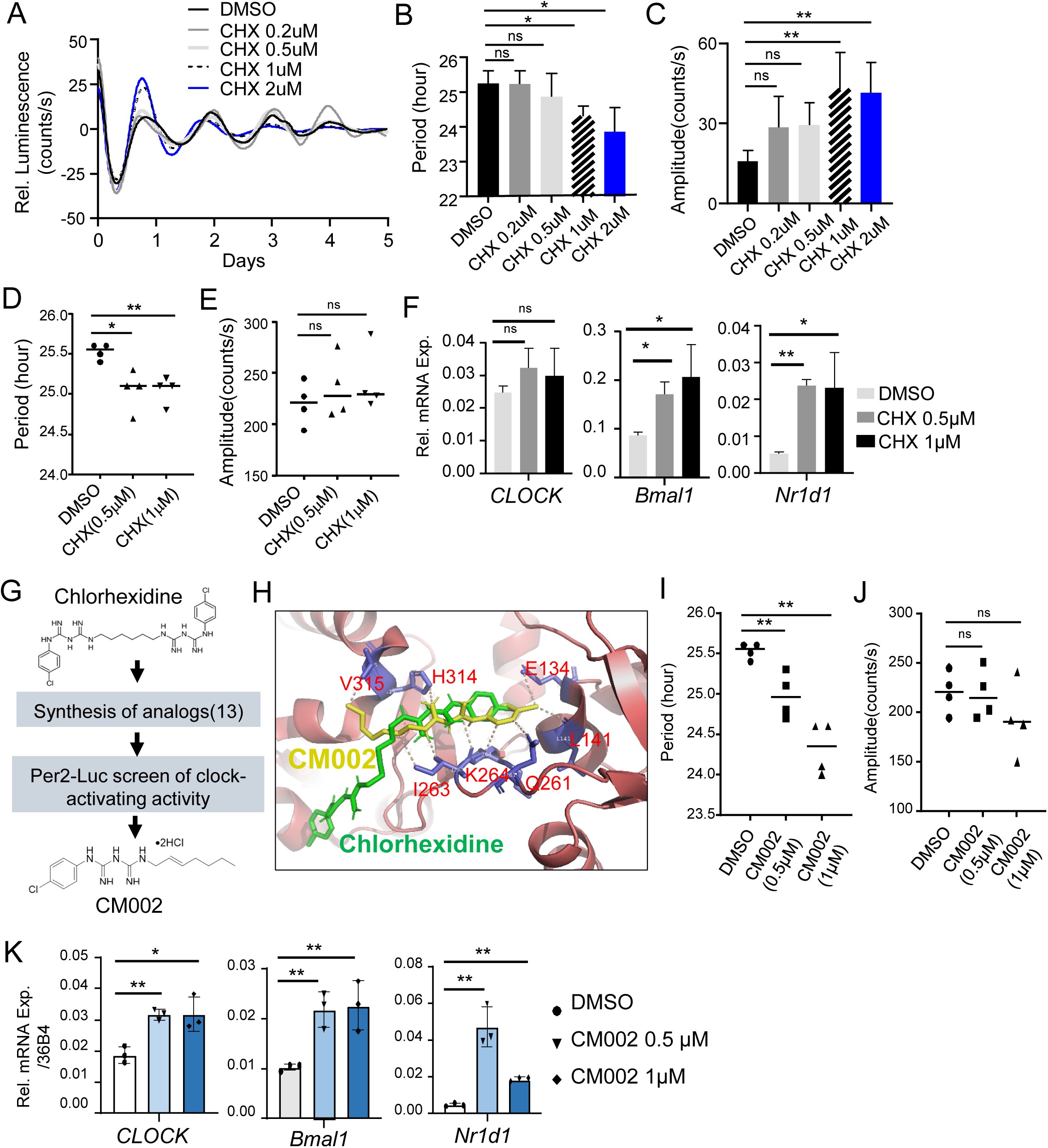
Clock-activating properties of chlorhexidine and a new derivative CM002 in adipogenic precursor cells. (A-E) The effect of chlorhexidine on modulating circadian clock activity. (A) Baseline-adjusted tracing plots of average luciferase bioluminescence activity of *Per2::dLuc* reporter-containing C3H10T1/2 mesenchymal stem cells, with quantitative analysis of clock period length (B) and cycling amplitude (C). (D, E) The effect of chlorhexidine on clock modulation in primary preadipocytes from *Per2::dLuc* knock-in mice, as shown by quantitative analysis of period length (D) and cycling amplitude (E) of luciferase reporter bioluminescence activity. CHX was added at indicated concentrations at the start of luciferase recording for 6 days (n=4). (F) RT-qPCR analysis of clock gene expression at indicated concentrations of CHX treatment for 6 hours in 3T3-L1 preadipocytes (n=3). (G) Schematic flowchart of lead compound discovery that identified CM002 as a new chlorhexidine analog with clock-activating properties. (H) Docking conformation of CM002 (yellow) within the CLOCK protein hydrophobic pocket together with chlorhexidine (green). Predicted CM002 interactions with CLOCK protein residues within 3-4A^0^ distance were shown. Crystal structure CLOCK (red) is based on PDB: 4f3l. (I, J) The effect of CM002 on clock modulation as shown by quantitative analysis of *Per2::dLuc* C3H10T1/2 reporter cells for clock period length (I) and cycling amplitude (J). CM002 at indicated concentrations were added at start of luciferase recording (n=4). (K) RT-qPCR analysis of clock gene expression of C3H10T/2 cells treated by CM002 at indicated concentrations for 6 hours (n=3). *, **: p<0.05 and 0.01 vs. DMSO by Student’s t test.

Based on the chemical scaffold of chlorhexidine, we generate 13 structural analogs via chemical synthesis through modification of the specific chemical groups. Using an *Per2::dLuc* luciferase-containing U2OS reporter line, screening of the clock-modulatory activities of these compounds led to identification of CM002, resembling approximately half of the CHX chemical structure (Fig. 1G & Fig. S2A), as a new clock activator with improved clock activity than that of CHX. CHX was identified through its structural fit for against a hydrophobic pocket of the PAS-A domain of the CLOCK protein [30]. We thus performed molecular docking analysis of CM002 in-silico binding mode within the shared hydrophobic pocket of CLOCK with CHX (Fig. 1H, S2A & S2B). Consistent with the overlap of CM002 structure with CHX scaffold, its predicted binding surface coincides with a large portion of CHX interactions within the CLOCK protein (Fig. 1H & S2B). A detailed examination of CLOCK protein residues with potential interactions with CM002 within 3-4A^0^ were illustrated (Fig. S2C). As shown by U2OS *Per2::dLuc* reporter, CM002 exhibited a dose-dependent effect on inducing clock period length shortening (Fig. 1I) without significantly affecting amplitude (Fig. 1G), demonstrating its clock-activating property. Furthermore, CM002 treatment of 10T1/2 cells led to induction of core clock genes, *CLOCK* and *Bmal1,* together with up-regulation of its direct target within the molecular clock loop, *Nr1d1* (Fig. 1K).

### 3.2 Chlorhexidine and CM002 inhibit adipogenic mesenchymal progenitor differentiation

Given that chlorhexidine and CM002 promote clock oscillation in adipogenic precursor cells, we determined whether these pharmacological clock activators are sufficient to inhibit adipocyte development. CHX treatment during adipogenic differentiation of 10T1/2 mesenchymal precursors led to a dose-dependent inhibition of adipocyte maturation, evident at the early stage of differentiation at 5 days following adipogenic induction, as indicated by phase-contrast, oil-red-O and Bodipy staining of lipid accumulation as a marker for mature adipocyte formation (Fig. 2A). The inhibitory effect of CHX on 10T1/2 differentiation remained at 9 days after differentiation (Fig. 2B & 2C). Further analysis of the adipogenic program demonstrated significantly impaired levels of adipogenic factors in day 8-differentiated C3H10T1/2 cells, including CEBP/α and PPARγ (Fig. 2D), together with attenuated expression of fatty acid synthase (FASN). The CHX analog CM002 displayed a stronger effect on inhibiting 10T1/2 adipogenesis, as indicated by its suppression of Bodipy-stained lipids that reached ∼50% inhibition at 1 µM (Fig. 2E). Consistent with its strong adipogenic-suppressive efficacy, this concentration of CM002 nearly abolished the expression of key adipogenic factors, CEBP/α and PPARγ, together with robust down-regulation of FABP4 and FASN (Fig. 2F). Thus, clock activation by CHX or CM002 was sufficient to suppress the lineage commitment and maturation of adipogenic mesenchymal precursor cells.

**Figure 2.**
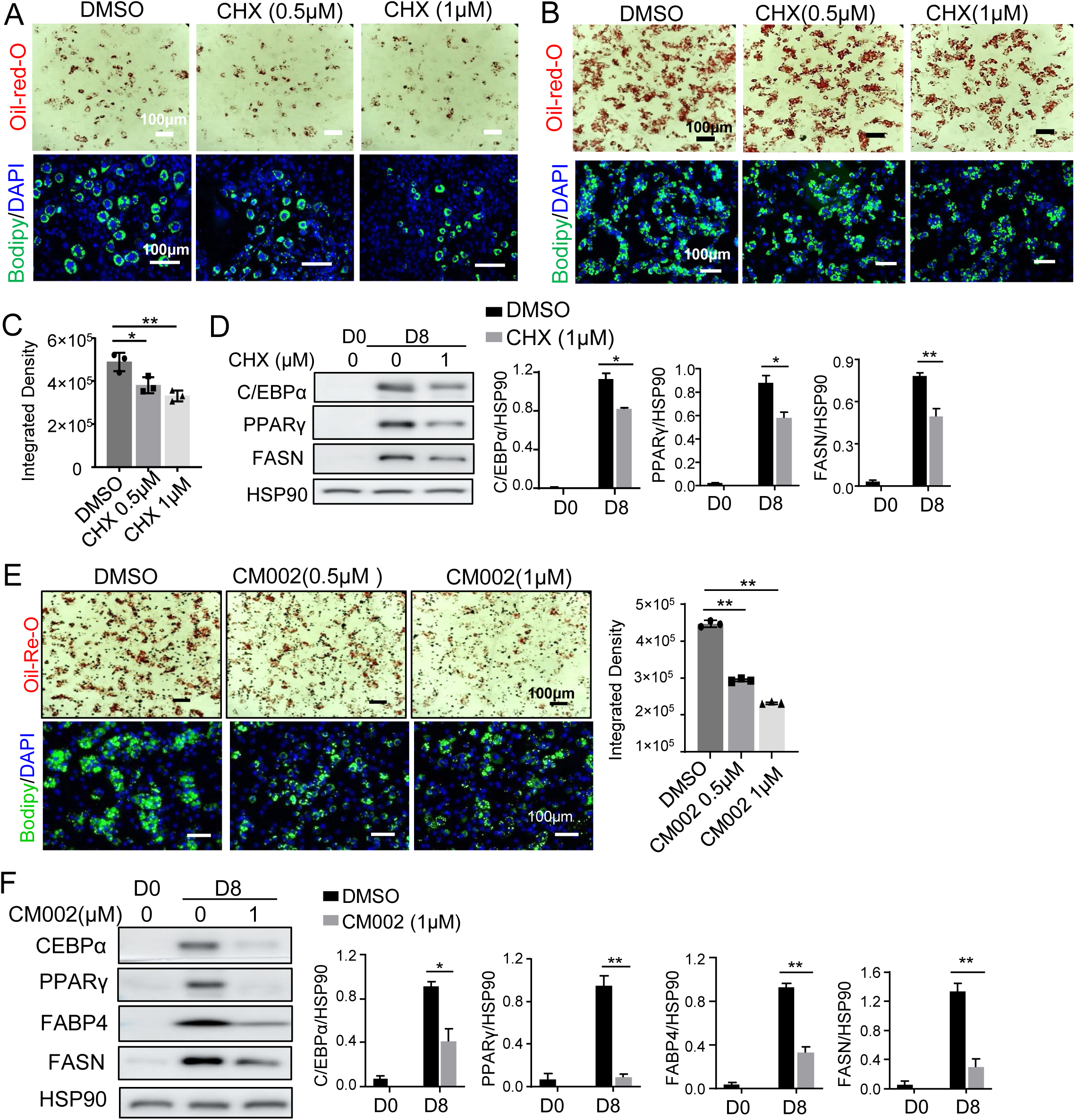
Inhibition of adipogenic mesenchymal precursor lineage commitment by chlorhexidine and CM002. (A-C) Representative images of oil-red-O and Bodipy staining of C3H10T1/2 cells at day 5 (A) and day 8 (B) of adipogenic differentiation treated with chlorhexidine at indicated concentrations, with quantification of Bodipy at day 8 (C). Average intensity of Bodipy fluorescence of three representative fields was obtained using Image J and normalized to DAPI signal. Scale bars: 100 μm. (D) Representative immunoblot analysis of adipogenic program before and after 8 days of adipogenic differentiation with 1μM of chlorhexidine treatment, with quantification of three replicates normalized to HSP90. (E) Representative images of oil-red-O and Bodipy staining of 10T1/2 cells after 8 days of adipogenic differentiation treated with CM002 at indicated concentrations, and quantitative analysis of Bodipy staining. (F) Representative immunoblot analysis of adipogenic program after 8 days of adipogenic differentiation treated with 1μM of CM002 and quantitative analysis. Each lane represents pooled sample of three replicates. *, **: p<0.05 and 0.01 vs. DMSO by Student’s t test.

### 3.3 Chlorhexidine and CM002 inhibition of terminal adipogenic differentiation of lineage-committed preadipocytes

We next determined whether pharmacological clock activation by CHX and CM002 could inhibit the terminal differentiation of lineage-committed preadipocytes. In 3T3-L1 preadipocytes, CHX was sufficient to reduce the formation of lipid-laden mature adipocytes in a dose-dependent manner, as indicated by oil-red-O staining and Bodipy staining following 6 days of adipogenic induction (Fig. 3A). This was further validated by quantitative analysis of Bodipy staining (Fig. 3B), and the attenuated levels of C/EBPα and PPARγ (Fig. 3C). CHX also displayed inhibitory effect on terminal differentiation of primary preadipocytes isolated from the stromal vascular fraction of adipose depot. This was evident by lipid staining at early differentiation day 4 (Fig. S3), or at day 6 of late differentiation with marked dose-dependent suppression of mature adipocyte formation (Fig. 3D-3E). At day 6 of differentiation, CHX did not attenuate protein expression of adipogenic factors (Fig. 3F). Instead, the marked reduction of FASN suggests its modulation of terminal differentiation through inhibition of lipid synthesis without significantly altering the adipogenic program. Using primary preadipocytes isolated from normal controls (FloxCtr) and clock-deficient Bmal1-null (BMKO) mice, we determined the effect of CM002 on adipogenic differentiation and whether its activity is dependent on clock modulation. CM002 treatment of normal controls led to dose-dependent inhibition of maturation, as indicated by lipid staining (Fig. 3G). In contrast, this effect on suppressing adipocyte formation was blunted in Bmal1-null preadipocytes, though not completely abolished. Similar effects were observed by oil-red-O staining (Fig. S4). Thus, CM002 displays inhibition of adipocyte terminal differentiation, and this effect was mediated, at least in part, by a clock-dependent mechanism.

**Figure 3.**
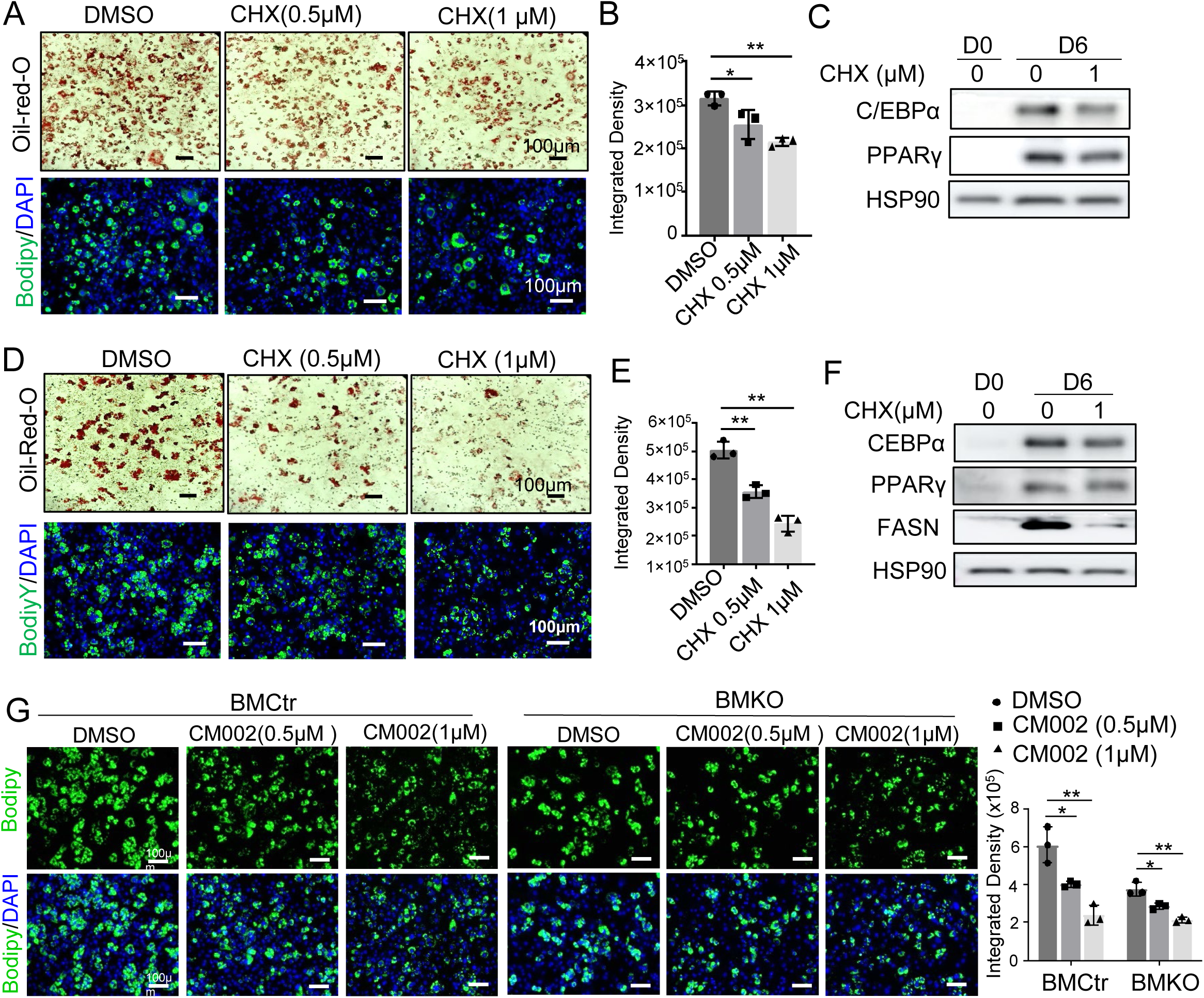
Inhibition of chlorhexidine and CM002 on terminal differentiation of preadipocytes. (A-C) Effect of CHX on 3T3-L1 preadipocyte differentiation. Representative images of oil-red-O staining and Bodipy fluorescence staining (A) with quantitative analysis of Bodipy staining of 3T3-L1 preadipocyte at day 6 of adipogenic differentiation with or without CHX at indicated concentrations (B), and immunoblot analysis of adipogenic factors (C). Scale bar: 100 μm. *, **: p<0.05 and 0.01 CHX vs. DMSO by Student’s t test. (D-F) Effect of CHX on terminal differentiation of primary preadipocytes from stromal vascular fraction of mice inguinal fat. Representative images of oil-red-O and Bodipy staining (D) with quantitative analysis of Bodipy at day 6 of differentiation with indicated CHX concentrations (E), and immunoblot analysis of adipogenic protein levels (F). (G) Effect of CM002 on inhibiting terminal differentiation of control (FloxCtr) or Bmal1-null (BMKO) primary preadipocytes. Representative images of Bodipy staining with CM002 treatment at indicated concentrations at day 6 differentiation were shown with quantitative analysis. Scale bar: 100 μm. *, **: p<0.05 or 0.01 CM002 vs. DMSO.

### 3.4 Chlorhexidine and CM002 induction of Wnt signaling in adipogenic precursors

The Wnt pathway is a strong inhibitory developmental signal of adipose tissue development and adipogenic differentiation [40], and circadian clock exerts direct transcriptional control of this signaling cascade to suppress adipogenesis. We thus determined whether the inhibitory effects of pharmacological clock activators on adipogenesis could be mediated by their modulation of CLOCK/Bmal1 target genes in Wnt signaling. In 3T3-L1 preadipocytes, we surveyed the effect of CHX on key Wnt signaling components that were identified as direct targets of Bmal1 [13, 34]. CHX displayed dose-dependent effects on inducing expression of up-stream Wnt ligands and receptors, *Wnt1*, *Wnt10a*, *Frizzled 5* (*Fzd5),* together with inductions of *Disheveled 2* (*Dvl2*) and *β-catenin* (Fig. 4A). Similar regulations were observed in 10T1/2 mesenchymal precursors (Fig. S5). During adipogenic differentiation, β-catenin protein level markedly declined from undifferentiated precursor state to mature adipocytes, as expected (Fig. 4B). Consistent with its effect on up-regulating β-catenin transcript, CHX was able to partially reverse the loss of β-catenin protein expression at day 8 of 10T1/2 adipogenic differentiation (Fig. 4B). Given that CHX induced the key components of Wnt signaling pathway, we further assayed Wnt signaling activity using a Wnt-responsive TOPFlash luciferase reporter containing TCF4 bindings sites under both basal and Wnt ligand-stimulated conditions [36]. Without ligand stimulation, CHX at 0.2 and 0.5 µM induced the luciferase reporter activity by ∼3 fold, and a similar degree of increased Wnt signaling by CHX treatment was observed upon addition of Wnt3a-containing media (Fig. 4C). In comparison, CM002 was also able to induce the up-regulation of Wnt ligands (Fig. 4D), whereas the ability of CM002 to re-activate β-catenin protein expression in day 6-differentiated adipocytes appeared stronger than that of chlorhexidine (Fig. 4E). Using the TOPFlash luciferase reporter, we directly compared the efficacy of CHX and CM002 to activate Wnt signaling. Under basal condition, CM002 was sufficient to stimulate Wnt activity to a similar degree as CHX (Fig. 4F). Upon Wnt3a stimulation, however, CM002 at lower concentrations of 0.1 and 0.2 µM induced TOPFlash activity comparable to that of CHX at 0.5 µM, suggesting improved Wnt-activating efficacy of CM002.

**Figure 4.**
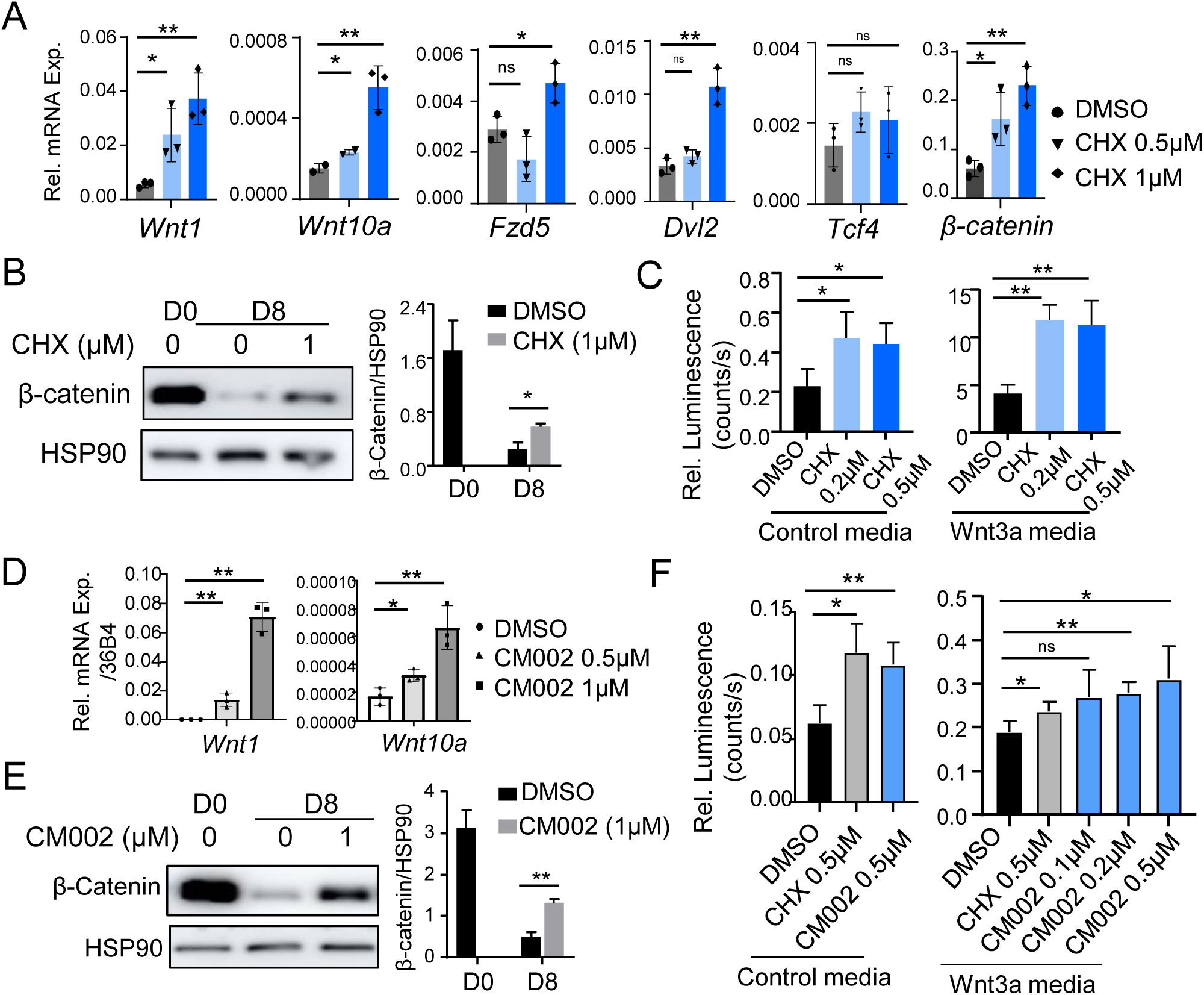
The effect of chlorhexidine and CM002 on promoting Wnt signaling in adipogenic progenitor cells. (A) RT-qPCR analysis of CHX effect on modulating expression of Wnt signaling pathway components in 3T3-L1 preadipocytes. Cells were treated with indicated concentrations of chlorhexidine for 6 hours. (B) Immunoblot analysis of CHX (1µM) on β-catenin protein level before and after C3H10T1/2 adipogenic differentiation for 8 days with quantitative analysis. (C) Analysis of chlorhexidine effect on Wnt signaling activity using a Wnt-responsive TOPFlash luciferase reporter assay under basal (control media) or Wnt3a-stimulated condition (10% Wnat3a-containing media). Data are presented as Mean ± SD of n=4 replicates. *, **: p<0.05 or 0.01 CHX vs. DMSO. (D) RT-qPCR analysis of CM002 effect on modulating Wnt ligand expression in 3T3-L1 preadipocytes at indicated concentrations for 6 hours. (E) Immunoblot analysis of CM002 effect on β-catenin protein level before and after C3H10T1/2 adipogenic differentiation for 8 days with quantitative analysis. (F) Comparison of chlorhexidine and CM002 on Wnt signaling activity using TOPFlash luciferase reporter. N=4 replicates. *, **: p<0.05 and 0.01 by ANOVA.

### 3.5 CM002 effect on reducing lipid storage in mature adipocytes

Adipocyte hypertrophy through excessive lipid storage is a major mechanism contributing to the development of obesity [41–43]. To explore its potential as for anti-obesity applications, we further determined whether CM002 modulates lipid remodeling by treating differentiated adipocytes for 2 days. CM002 treatment of adipocytes differentiated from primary mouse preadipocytes resulted in marked lowering of the amount of lipids as indicated by Bodipy staining (Fig. 5A). This was accompanied by the reduction of a key enzyme in fatty acid synthesis (FASN) and the fatty acid transport protein (FABP4, Fig. 5B), suggesting concerted mechanisms in suppressing lipid accumulation in adipocytes by CM002. CM002 was able to induce clock gene expression, including *CLOCK*, *Dbp* and *Nr1d2,* indicative of clock activation in differentiated adipocytes (Fig. 5C). In addition, expression analysis revealed up-regulation of thermogenic regulator uncoupling protein 1 (*Ucp-1*) and the lipolytic enzyme hormone-sensitive lipase (*HSL*), suggesting potential browning effect that may enhance lipid remodeling in adipocytes (Fig. 5D). Using differentiated *Bmal1-null* primary adipocytes, we tested the clock-dependency of CM002 inhibitory effect on adipocyte lipid storage. As expected, CM002 significantly reduced lipid accumulation in adipocytes derived from floxed control mice, whereas in adipocytes with Bmal1 ablation, this effect was completely lost (Fig. 5E). Thus, in addition to inhibiting adipocyte development, CM002 also limits fat storage in adipocytes that may prevent hypertrophy.

**Figure 5.**
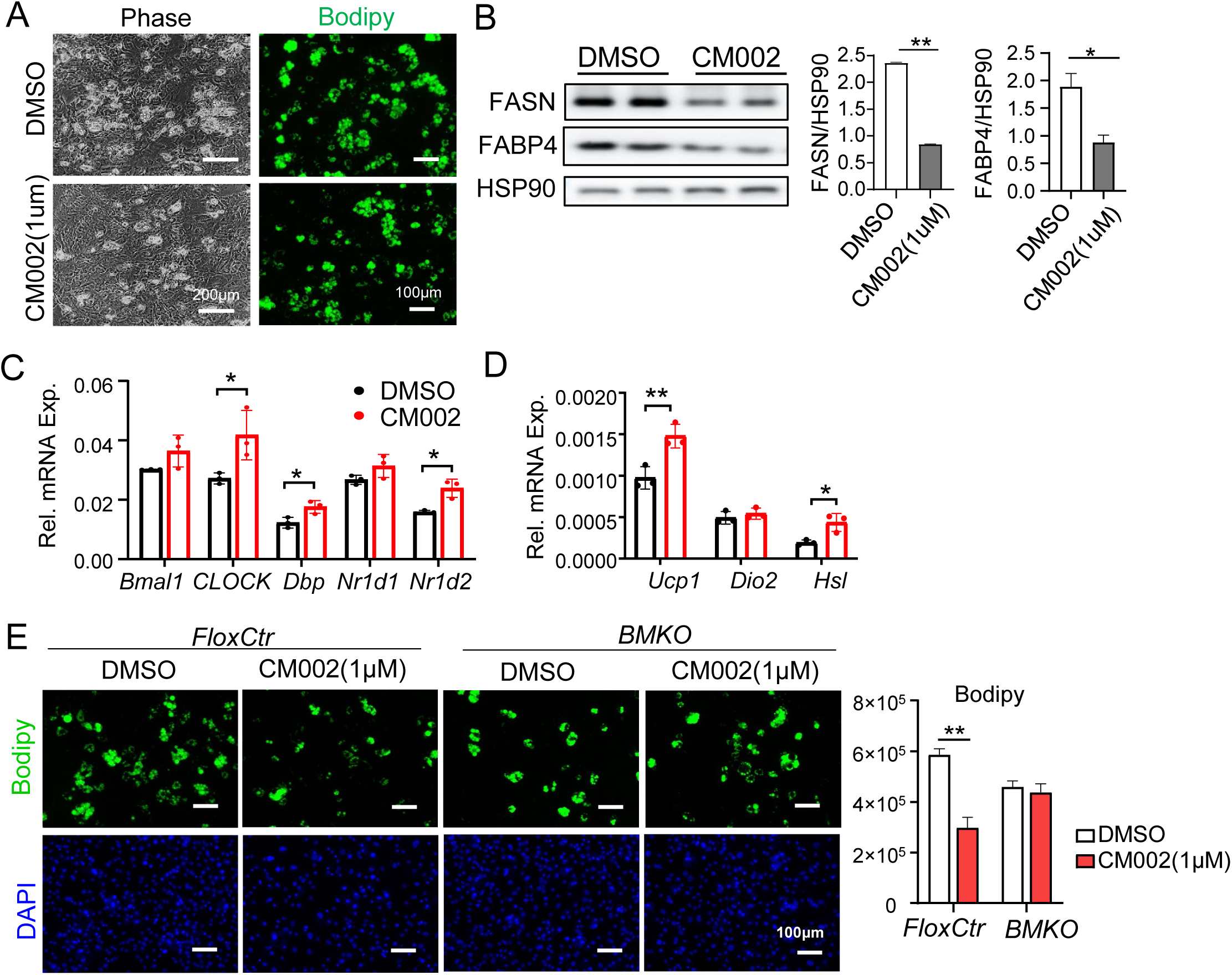
CM002 inhibition of lipid storage in mature adipocytes. (A-D) Primary mouse preadipocytes were subjected to adipogenic differentiation for 4 days prior to treatment with CM002 for 48 hours. Representative phase-contrast images of adipocyte morphology and Bodipy staining treatment (A), immunoblot analysis (B), and RT-qPCR analysis of clock (C) and thermogenic genes (D) in adipocytes with or without 1µM of CM002 treatment. (E) Representative images of Bodipy staining of control (FloxCtr) or Bmal1-null (BMKO) adipocytes treated with 1µM CM002 for 48 hours with quantitative analysis. Scale bar: 100 μm. *, **: p<0.05 or 0.01 CM002 vs. DMSO by Student’s t test.

To explore the translational potential of CM002, we used primary preadipocytes derived from visceral fat depots of human donors to further examine its modulation of adipogenesis and lipid storage in mature adipocytes. When human preadipocytes were treated with CM002 throughout the differentiation time course, there was a robust effect on suppressing adipogenic maturation as compared to DMSO control (Fig. 6A). Furthermore, CM002 displayed similar inhibition of lipid deposition in late-stage differentiated human adipocytes following adipogenic induction for 6 days (Fig. 6B). This was accompanied with strong induction of clock proteins BMAL1 and RORα (Fig. 6C), suggesting CM002 activation of clock in human adipocytes, together with inhibition of proteins involved in lipid synthesis and transport (Fig. 6D).

**Figure 6.**
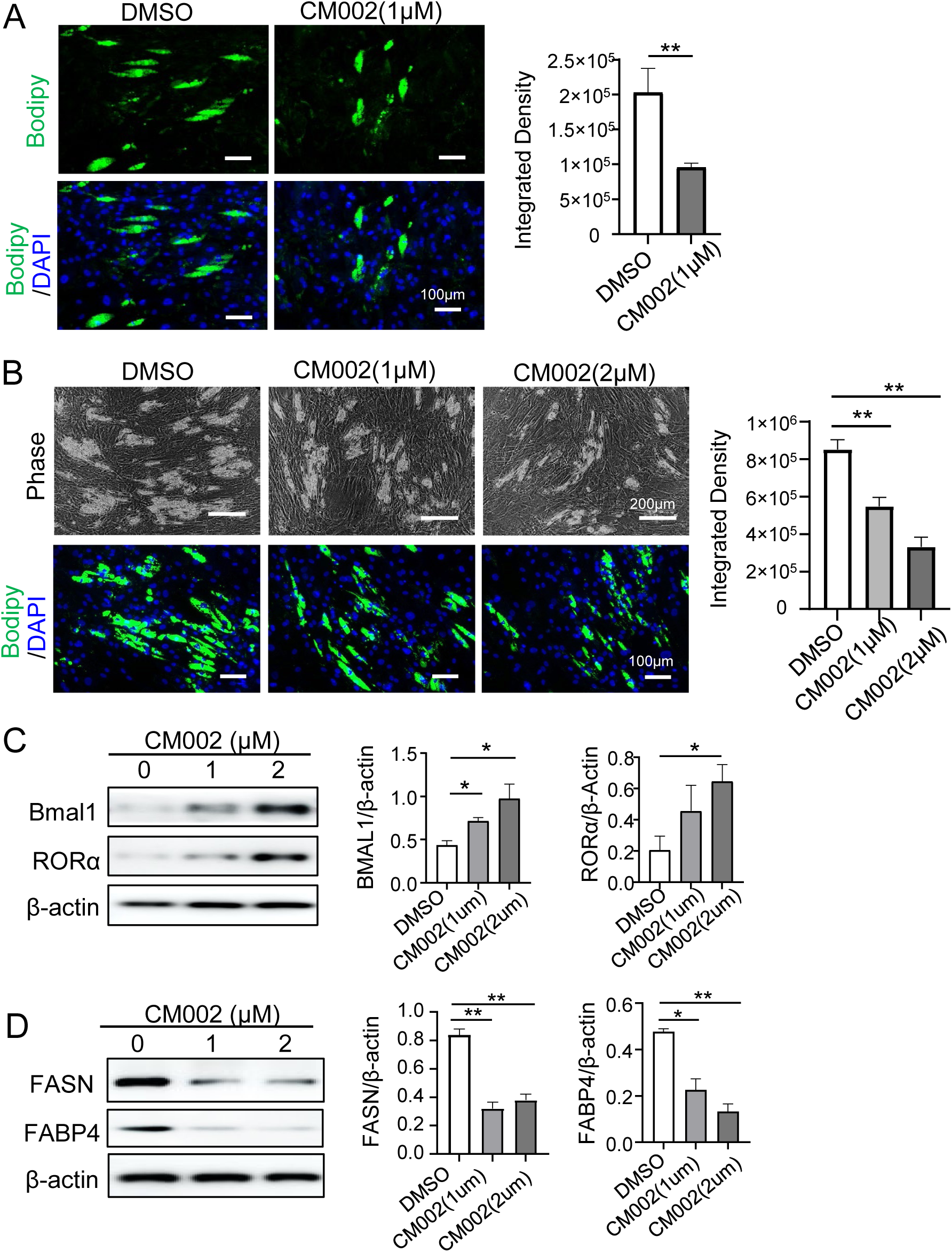
CM002 suppression of adipogenic differentiation and lipid storage in human adipocytes. Human primary preadipocytes were obtained from subcutaneous fat and expanded in vitro for two passages followed by adipogenic differentiation for 10-12 days. (A) Representative images of Bodipy staining of differentiated human adipocytes treated with 1µM CM002 through the differentiation time course with quantitative analysis. (B) Representative images of phase-contrast and Bodipy staining of 12 day-differentiated human adipocytes with CM002 treatment at indicated concentrations for last 4 days of differentiation. Quantitative analysis of lipid staining was shown in right panel. (C, D) Immunoblot analysis of CM002 effect on clock proteins (C), and proteins involved in lipid remodeling (D) in human adipocytes. Each lane represents pooled sample of three replicates. *, **: p<0.05 or 0.01 CM002 vs. DMSO by Student’s t test.

### 3.6 CM002 induced clock activation with reduction of fat mass in mice on regular chow diet

The robust CM002 effects on suppressing the development and fat storage of adipocytes suggest its potential for anti-obesity applications. We thus determined its *in vivo* efficacy on body weight and adipose tissues under chow and high fat diet-feeding conditions. Initial determination of the maximal lethal dose in wild-type C57/BL6 mice and pharmacokinetics study revealed that 5-10 mg/Kg daily IP dose were suitable for *in vivo* testing, without overt liver toxicity detected for 5mg/Kg dosing as shown by serum ALT analysis of mice treated with CM002 for 15 days on chow diet (Fig. S6A) or for 7 days on a high-fat diet (Fig. S6B). IP delivery of CM002 at 5mg/Kg was sufficient to induce clock protein induction in adipose tissue, as shown for BMAL1, RORα and DBP (Fig. 7A). Similar effects on clock activation were found in skeletal muscle (Fig. S6C). Treatment of CM002 for 18 days in 24-week-old wild-type mice led to significant reductions of body weight of up to ∼15%, as compared with vehicle-treated controls that maintained their weight for this duration (Fig. 7B). The reduction of body weight is mostly due to the loss of fat mass (Fig. 7C), whereas total lean mass of CM002-treated mice remained similar to before treatment with increased relative percentage due to predominant reductio of fat (Fig. 7D). Histological examination of distinct adipose depots, the visceral epididymal white adipose tissue (eWAT), subcutaneous inguinal adipose tissue (iWAT), revealed markedly attenuated adipocyte hypertrophy, with reduced lipid storage in interscapular brown adipose tissue (BAT, Fig. 7E). Further analysis of protein expression in eWAT white adipocyte depot revealed significantly markedly lower levels of adipogenic factors C/EBPα and PPARγ together with the master transcription regulator of lipogenesis SREBP-1c and fatty acid synthetase (FASN) (Fig. 7F). Interestingly, the overall energy expenditure of CM002-treated animals was comparable to that of controls, as indicated by the level of oxygen consumption (VO2, Fig. 7G). However, there was a notable reduction of respiratory exchange ratio (RER) during both the light and dark cycles, suggesting a preferential utilization toward fat oxidation over carbohydrate (Fig. 7H). Food intake and total ambulatory activity levels, and their day/night diurnal rhythms, are not altered in CM002-teated cohort (Fig. 7I & 7J). Consistent with its efficacy in reducing fat mass and RER, CM002 administration significantly reduced circulating free fatty acid levels in plasma (Fig. 7K). In addition, fasting glucose level was also lower in the CM002-treated cohort (Fig. 7L), and glucose tolerance test further revealed enhanced glucose disposal in these mice (Fig. 7M). In skeletal muscle, the insulin-responsive glucose transporter GLUT4 protein was elevated, along with a protein expression profile indicative of enhanced mitochondrial abundance or activity as shown by increased PGC-1α, succinate dehydrogenase (SDHB) and Tom 20 (Fig. 7N). Thus, CM002 displays clock-enhancing properties in vivo, and reduced lipogenesis in adipose tissue coupled with increased metabolic oxidation in muscle may underly its anti-obesity and glucose-lowering effects.

**Figure 7.**
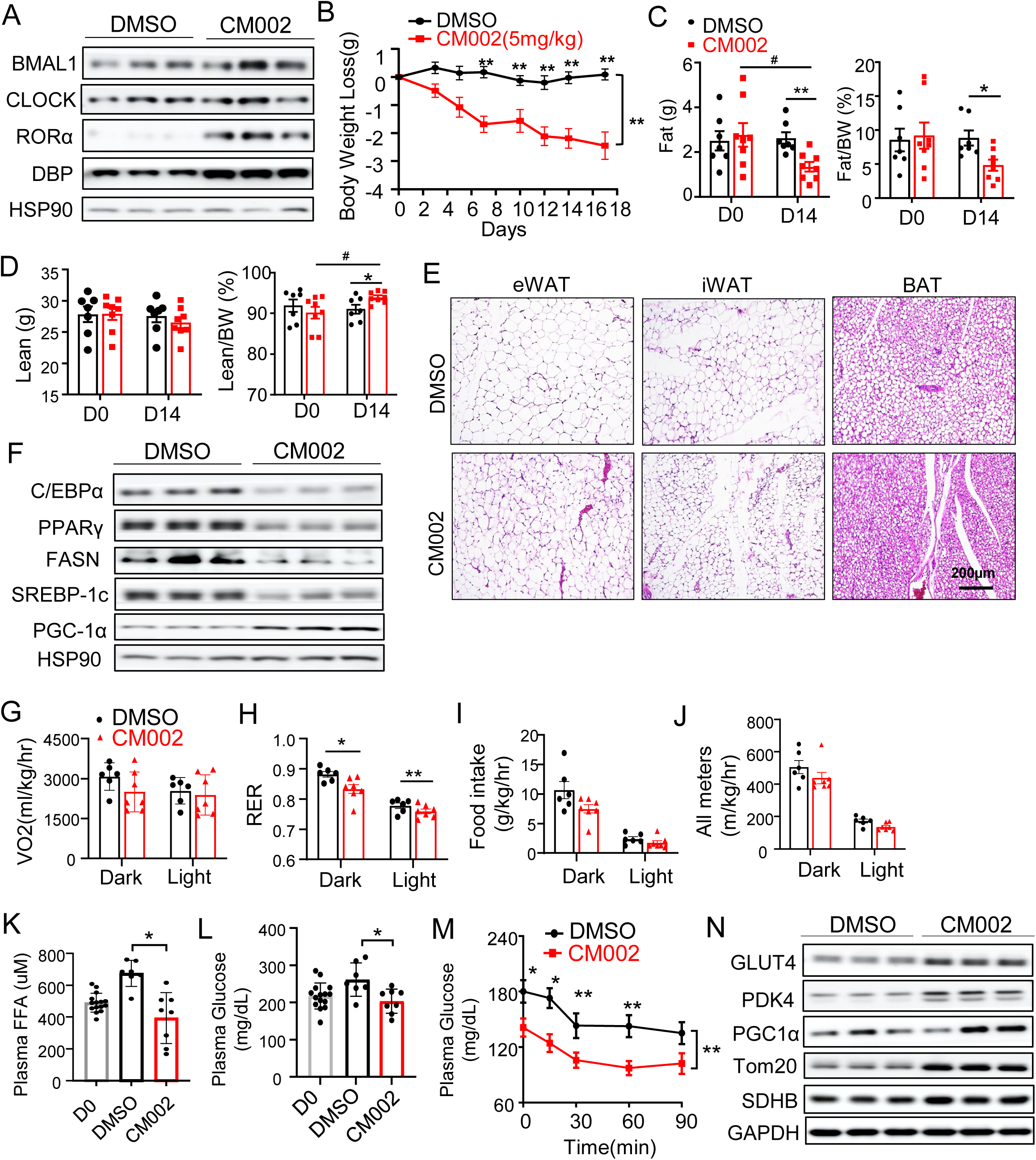
Reduced fat mass by CM002 treatment in mice on regular chow diet. (A) Immunoblot analysis of clock proteins in visceral fat of C57/BL6 wild-type mice on regular chow diet treated for 18 days with vehicle control (0.1% DMSO) or CM002 (5mg/Kg). Each lane represents a pooled sample of 2-3 mice. (B) Loss of body weight of mice during CM002 treatment for 18 days. Mice were treated three times per week with 5mg/kg of CM002 via intraperitoneal injection. (n=7-8/group). **: P≤0.05 one way ANOVA with Tukey’s post-hoc analysis. (C, D) NMR analysis of total fat content (C) and lean mass (D) with relative percentage at the start and after 14 days of CM002 treatment. (E) Representative images of H/E histology of epidydimal white adipose tissue (eWAT), inguinal white adipose tissue (iWAT), and interscapular brown adipose tissue (BAT) in DMSO or CM002-treated mice after 18 days of CM002 treatment. (F) Immunoblot analysis of adipogenic and lipogenic proteins in iWAT of DMSO or CM002-treated mice. (G-J) Analysis of energy balance by indirect calorimetry in mice on chow diet treated with CM002 for 7 days, as indicated by oxygen consumption rate (VO2, G), respiratory exchange ratio (RER, H), food intake (I) and ambulatory activity (J). (n=7-8/group). *, **: p<0.05 or 0.01 CM002 vs. DMSO (K, L) Plasma levels of free fatty acid (FFA, K) and glucose (L) in DMSO or CM002-treated mice after 18 days. (M) Insulin tolerance test in DMSO or CM002-treated mice after 18 days of treatment. *, **: p<0.05 or 0.01 one-way ANOVA with Tukey’s post-hoc analysis (n=7-8/group). (N) Representative images of immunoblot analysis of proteins involved in muscle metabolism in DMSO or CM002-treated mice. Each lane represents a pooled sample of 2-3 mice.

### 3.7 Anti-obesity efficacy of CM002 treatment in mice on high-fat diet feeding

We next determined the potential for CM002 administration to counter the development of obesity in mice with high-fat-induced obesity with testing of distinct dosing regimens. Initial study of daily CM002 delivery at 5mg/Kg led to marked loss of up to 8 grams of body weight in the high-fat diet mice cohort within a week (Fig. 8A), DMSO-treated controls also lost weight, though significantly less in comparison with CM002. The reduction of body weight in CM002-treated mice was largely due to fat loss (Fig. 8B), although with moderate loss of lean mass in both DMSO and CM002-treated groups (Fig. S6D). Interestingly, though both visceral and inguinal fat depots (Fig. 8C & 8D) in CM002-treated mice displayed tendency toward lower fat mass, only brown fat weight was significantly reduced (Fig. 8E). H/E histology demonstrated marked reduction of lipid accumulation in all adipose depots examined (Fig. 8F). In line with these findings, CM002 reduced the protein expression of adipogenic factors and lipogenic enzymes in inguinal and brown adipose depots (Fig. 8G & 8H). Analysis of clock proteins confirmed the inductions of BMAL1, CLOCK and RORα in these tissues (Fig. S6E & S6F). CM002-treated HFD-fed mice displayed significantly lower circulating free fatty acids levels (Fig. 8I), though only with a tendency toward lower plasma glucose (Fig. 8J). Interestingly, serum triglyceride levels were not altered by CM002 (Fig. 8K). As the initial DMSO-treated groups displayed significant loss of body weight, considering these findings, we further tested whether a less frequent IP dosing regimen with longer duration could be better tolerated for mice on high-fat diet feeding with potential for sustained anti-obesity efficacy. Using three 7mg/Kg doses per week for up to 16 days, this CM002 regimen recapitulated findings from daily treatment. A similar amount of weight reduction of ∼8g was achieved in this CM002-treated cohort at 16 days (Fig. 8L & 8M). In contrast to earlier treatment, body weight of the vehicle-treated mice was largely maintained with less frequent injections. The weight loss induced by CM002 under this dosing regimen could be solely accounted for by the loss of fat mass, while total lean mass was not significantly altered as indicated by NMR analysis of body composition before and after 14 days of CM002 treatment (Fig. 8N). Dissection of individual fat depots further corroborated the reduced wet weight of distinct depots examined, including eWAT, iWAT and BAT (Fig. 8O-8Q). In contrast, individual muscle weight examined remained comparable to that of vehicle control cohort, such as shown for the Tibialis Anterior (Fig. 8R). Collectively, our in vivo findings of CM002 treatment under chow or diet-induced obesity conditions revealed its consistent efficacy on inducing loss of fat mass with associated insulin-sensitizing effects.

**Figure 8.**
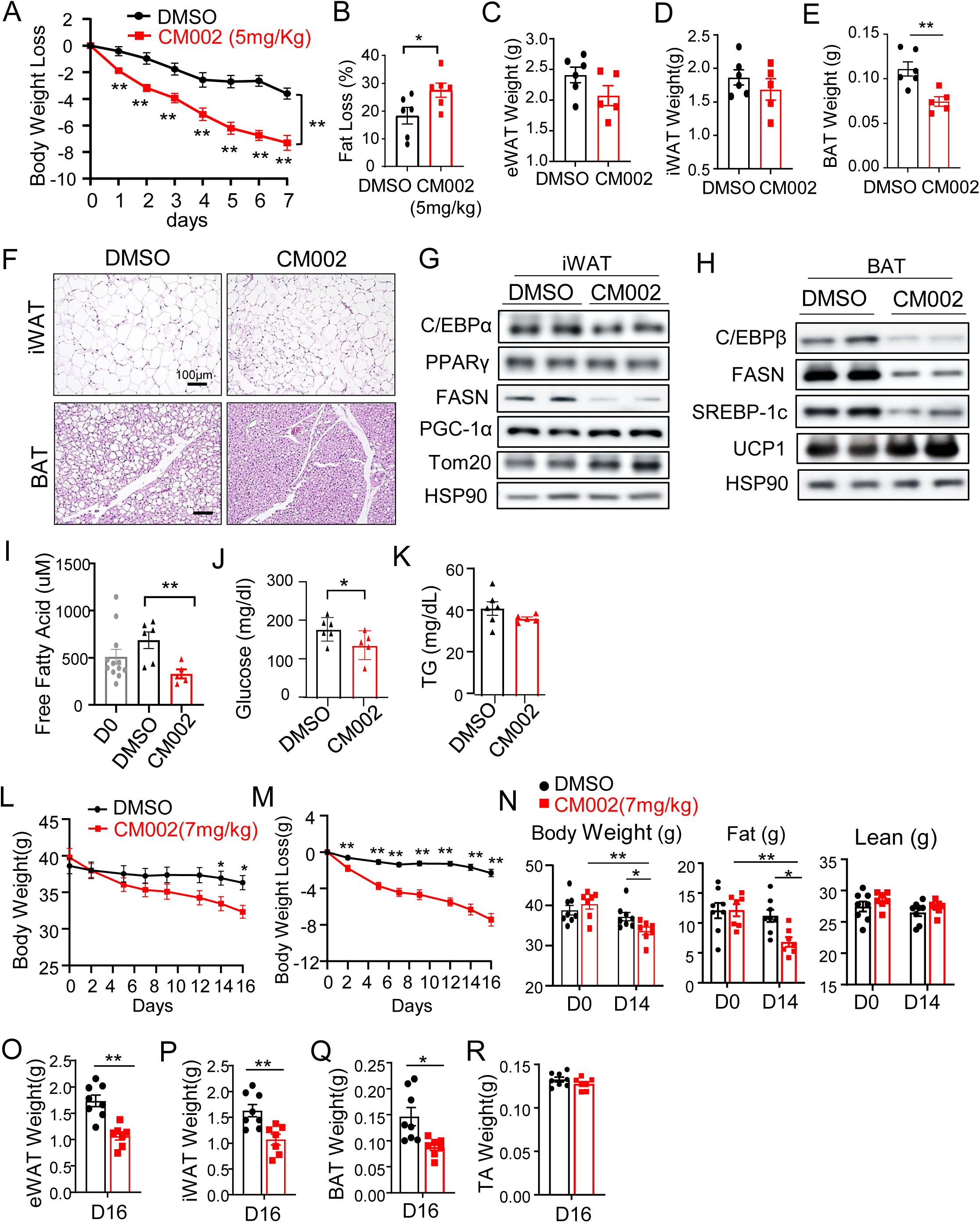
Anti-obesity effect of CM002 treatment in mice with high-fat diet feeding. (A) Analysis of loss of body weight of mice on high-fat diet with daily intraperitoneal injection of 5mg/kg of CM002 for 7 days. **: P≤0.05 one way ANOVA with Tukey’s post-hoc (n=5-6/group). (B) NMR analysis of total fat content after 7 days of treatment plotted as change from before CM002 injection. (C-D) Analysis of dissected tissue weight of eWAT (C), iWAT (D), and BAT (E) after 7 days of CM002 treatment. (F) Representative images of H/E histology of iWAT and BAT in DMSO or CM002-treated mice after 7 days. (G, H) Immunoblot analysis of adipogenic and lipogenic protein expression in iWAT (G) and BAT (H) of control and CM002-treated mice. Each lane represents a pooled sample of 2 mice. (I-K) Fasting plasma levels of FFA (I), glucose (J) and triglyceride (K) in control mice or after 7 days CM002 treatment. (L-R) Effect of three times weekly CM002 treatment at 7mg/Kg for 16 days on body weight and tissue mass. (n=7-8/group). Analysis of absolute body weight (L) and change from baseline in each group (M) in mice treated by vehicle or CM002. **: P≤0.05 one way ANOVA with Tukey’s post-hoc. (N) NMR analysis of total fat mass and lean mass after 14 days of vehicle or CM002 treatment. (O-R) Analysis of dissected tissue weight of eWAT (O), iWAT (P) and BAT (Q) in mice treated with vehicle or CM002 for 16 days. *, **: P≤0.05 or 0.01 by Student’s t test.

## 4. DISCUSSION

Circadian clock plays a key regulatory role in adipogenesis, and a large body of evidence indicates that clock disruption leads to the development of obesity and insulin resistance [3]. In light of the widespread “social jetlag” in a modern lifestyle, circadian clock could be a potential therapeutic target to counter the current epidemic of metabolic diseases [19, 29]. Employing distinct cellular models, our current study uncovered the anti-adipogenic properties of novel clock-activating compounds, CHX and CM002. Furthermore, findings of the *in vivo* anti-obesity efficacy of CM002 demonstrated its potential toward clock-targeted drug development for metabolic disease therapy.

Studies to date have established a strong association between circadian clock disruption and the development of metabolic disorders [3, 44]. Given the pervasive temporal regulation of key metabolic pathways and the daily oscillations of metabolites, dysregulation of clock-controlled metabolic outputs due to circadian misalignment collectively impacts metabolic homeostasis. We previously reported that clock transcription factor Bmal1 exert direct transcriptional control of Wnt signaling pathway components leading to inhibition of adipogenesis [13]. Loss of clock function in mice, due to deficiency of core clock transcription activators CLOCK or Bmal1, leads to obesity [13–15]. In addition, disruption of clock via various environmental clock manipulations predispose to the development of obesity [7, 45–47], while shiftwork poses a significant risk for insulin resistance and type II diabetes [4, 8, 47]. Conversely, maintaining or re-enforcing clock function provide protection against metabolic dysfunctions, particularly due to circadian misalignment. Nobiletin, a clock amplitude-enhancing molecule that was identified as an agonist for the positive clock regulator RORα [25], was found to directly suppress adipogenesis with demonstrated anti-obesity effect [26]. Along this line, a structurally related flavonoid molecule naringenin with RORα agonist activity also displayed significant effect on inhibiting adipogenic maturation while inducing marked loss of fat mass in mice [48].

We thus first explored the anti-adipogenic potential of chlorhexidine, a clock-activating molecule recently identified from a HTS screen for clock modulators [30]. Further chemical derivation of chlorhexidine led to the identification of CM002 with improved clock-modulatory activity. In line with their activation of the Wnt signaling that mediates clock inhibition of adipogenesis [13], these clock modulators were able to inhibit distinct stages of adipocyte development involving commitment of mesenchymal stem cell and terminal differentiation of lineage-determined preadipocytes. These initial findings support the notion that pharmacological interventions to promote clock function can inhibit the maturation of adipogenic progenitors that could be applicable for obesity treatment. Notably, CM002 also induced clock activation in differentiated adipocytes, limiting lipid storage with suppression of adipogenic factors and lipogenic enzymes indicative of its potential to negatively impact adipocyte hypertrophy. Furthermore, these effects of CM002 on inhibiting adipogenic maturation were recapitulated in human adipocytes. Corroborating these *in vitro* findings of CM002 inhibition of adipocyte maturation and hypertrophy-associated fat accumulation, CM002 treatment led to robust anti-obesity effect with marked loss of fat mass, under either chow or high-fat diet conditions. Collectively, these results provided direct support for pharmacological targeting of clock function to suppress the adipogenic and lipogenic drive of adipocytes as a novel avenue for anti-obesity therapy. As small molecule clock activators likely induce clock function in distinct tissues, additional mechanisms could be attributable to the overall metabolic benefits that warrant investigations.

Medicinal chemical optimization of the chlorhexidine scaffold led to the discovery of CM002. In addition to its improved efficacy range on inducing clock period shortening without cellular toxicity within the concentrations tested, its stimulation of Wnt signaling was improved as compared to chlorhexidine as well as inhibition of the adipogenic program. Our observation of the differential modulation of CHX and CM002 on adipogenic genes between mesenchymal precursor and lineage-committed progenitors, C3H10T1/2 cells as compared to 3T3-L1 and primary preadipocytes, is intriguing. Both CHX and CM002 markedly attenuated adipogenic differentiation of mesenchymal precursor cells, accompanied with significant downregulation of key adipogenic factors and mature adipocyte markers. Interestingly, in preadipocytes, despite comparable inhibition of adipocyte maturation as indicated by lipid accumulation, their effects on adipgenic program were confined to blocking mature adipocyte marker expression without significantly altering adipogenic factors. These results suggest potential developmental stage-specific regulations of the differentiation process by these clock activators. The ability of CM002 to limit lipid accumulation in differentiated adipocytes in a clock-dependent manner, and its effectiveness in human adipocytes, corroborates a key mechanism of action that could be most relevant to prevent adipocyte hypertrophy in the development of obesity.

High-fat diet feeding-induced obesity leads to dampening of clock oscillation, raising the possibility that re-enforcing clock function may reverse the adverse effect of nutritional overload on clock-controlled metabolic pathways [21]. *In vivo* studies of CM002 in the high-fat diet-induced obesity model revealed its robust effect on preventing adipose tissue expansion. This anti-obesity effect of CM002 was reproducible when tested using different dosing regimens with less frequent injections to avoid adverse influence on lean mass. The normal range of serum ALT levels in CM002-treated animals demonstrated the absence of hepatic toxicity. In addition, food intake and activity levels of these mice were comparable to that of controls, with the day-night diurnal rhythm maintained. The effect of CM002 on reducing fat mass was evident in all fat depots examined. As we demonstrated under both chow and high-fat diet conditions, these effects on adipose tissues were mediated, at least in part, by inhibition of the lipogenic and adipogenic programs. Interestingly, up-regulation of mitochondrial genes and elevated expression of UCP-1 were found in beige and brown adipose tissues, implicating potential browning effect that may promote fat oxidation in thermogenic fat depots. CM002 induced clock activation in adipose tissues, while similar effects are also observed in skeletal muscle and liver (data not shown). Thus, it is possible that the systemic metabolic impacts of CM002, including reduction of circulating free fatty acids and glucose levels, could be result of the collective contribution of beneficial effects in distinct metabolic sites, in addition to its anti-adipogenic and anti-lipogenic actions. Interestingly, the effect of CM002 on attenuating the glucose excursion as revealed by insulin tolerance test could be mostly accounted for by the lowered fasting glucose level, suggesting potential suppression of glucose production from the liver. Nonetheless, the elevated GLUT4 protein level in skeletal muscle, together with inductions of PGC-1α and SDHB, indicate that improved muscle metabolism may also contribute to the systemic metabolic benefits of CM002 treatment. Considering the known metabolic functions of circadian clock in tissues that are integral to metabolic homeostasis [37, 49–53], additional effects of CM002, in addition to adipose tissue regulations, warrant further detailed investigations.

It is conceivable the further optimization of CM002 scaffold is required to obtain more potent clock-activating compounds with improved efficacy for anti-obesity applications. Clock exerts anti-adipogenic effect while promoting myogenesis [13, 30, 34], and BMAL1 is directly involved in stimulating protein translation downstream to mTOR signaling in the ribosome [54]. Notably, for in vivo administrations of CM002 with its strong effect on inducing ∼40% loss of fat mass in the diet-induced obesity model, it largely maintained the muscle mass similar to that of the control cohorts. The potential muscle-preserving properties of CM002, or with its more potent analogs, could be further examined in appropriate models. In addition, clock function is dampened during aging with loss of cycling amplitude and a switch toward stress response [55, 56], while re-enforcing clock modulation may lead to anti-aging benefits [57–59]. Discovery of diverse chemical modulators to augmenting clock may provide protection against aging-associated metabolic dysfunctions or pathologies. Given the growing attention to target the circadian clock for disease interventions, pharmacological means to re-enforce endogenous temporal control of biological processes may lead to therapeutic development in a diverse array of human pathologies associated with circadian misalignment.

In summary, our study uncovered, for the first time, the anti-adipogenic properties of new clock-modulating compounds that hold potential for anti-obesity drug development. With the widespread metabolic consequences of circadian clock disruption, development of clock-activating molecules to maintain metabolic homeostasis may offer new avenues for the prevention or treatment of obesity and related metabolic consequences.

## Acknowledgements

We thank Drs. Steven Kay at University of Southern California for sharing luciferase reporter cell lines used in this study, and Drs. Seung-Hee Yoo and Zheng Chen at University of Texas at Houston Health Science Center for providing the Per2-luciferase plasmids. We thank the City of Hope Shared Resources Animal Phenotyping for carrying out metabolic phenotyping analysis. KM is a faculty member supported by the NCI-designated Comprehensive Cancer Center at the City of Hope National Cancer Center. This project was supported by National Institute of Health grants R01DK112794, R56AG080294, and an Innovative Pilot Award from Arthur Riggs Diabetes and Metabolism Research Institute to KM. The funders had no role in study design, data collection and analysis, decision to publish, or preparation of the manuscript.

## CRediT Author contributions statement

XX, JP, TK and LT: data curation and investigation, methodology, formal analysis, manuscript editing; ZF, AA WH and DH: Software, data curation, manuscript review and editing; KM: formal analysis, project administration, manuscript writing and editing, and funding acquisition.

## Ethics approval and consent to participate

All animal experiments in this study were approved by the Institutional Animal Care & Use Committee (IACUC) of City of Hope according to the approval. The protocol number: 17110, entitled “circadian regulation of metabolism” with approval date is from 12/11/2020 to 12/10/2023.

## Data Availability

All data generated and analyzed during this study are included in this published article and associated Supplementary Information files.

