## Supplementary material for "Novel circadian clock activators display anti-obesity efficacy via suppression of adipocyte development and hypertrophy": Suppl data

#### Supplemental Figure S1.

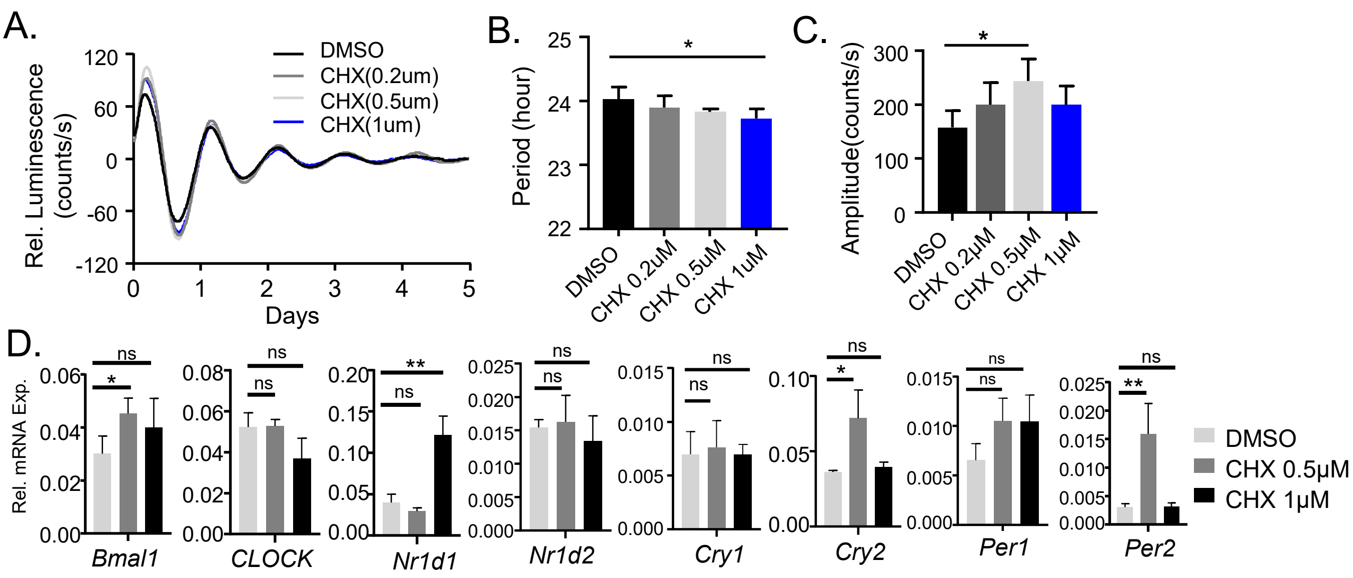


**Figure S1. Effect of chlorhexidine as a clock activator in 3T3-L1 preadipocytes.** (A) Baseline-adjusted tracing plots of average bioluminescence activity of Per2::dLuc reporter-containing 3T3-L1 preadipocytes (D-F) for 5 days, with quantitative analysis of clock period length (B) and cycling amplitude (C). Chlorhexidine (CHX) treatment was added to culture media at indicated concentrations. Data are presented as Mean ± SD of n=4 replicates for each concentration tested, for three independent repeat experiments. (D) RT-qPCR analysis of clock gene expression at indicated concentrations of CHX treatment for 6 hours in C3H10T/2 cells. Data are presented as Mean ± SD of n=3 replicates. *, **: p<0.05 and 0.01 CHX vs. DMSO by Student's t test.

#### Supplemental Figure S2.

A.

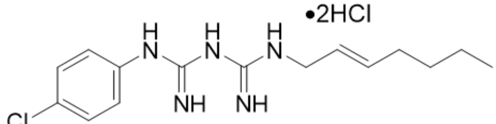


CM002

Chemical Formula: $C_{15}H_{24}Cl_3N_5$
Molecular Weight: 380.7420

B.

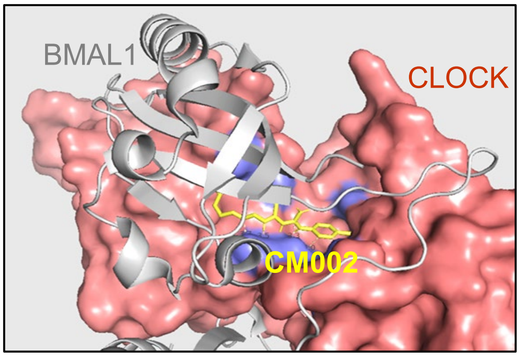


C.

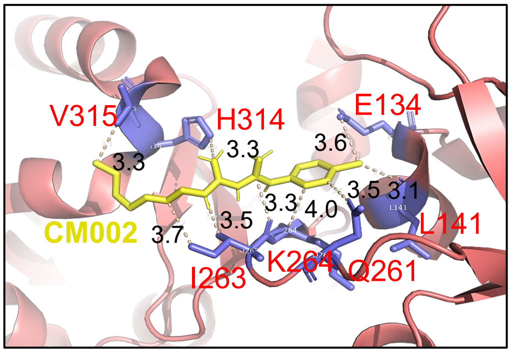


**Figure S2.** (A) Chemical structure of CM002. (B) Molecular docking modeling of CM002 (yellow) structure together with hydrophobic interaction pocket between CLOCK (pink) and Bmal1 protein (gray) PAS-A domain. Crystal structure shown for CLOCK (red, surface mode) with BMAL1 (grey, cartoon mode) is based on PDB: 4f3l. (C) Predicted CM002 interactions with CLOCK protein residues within 3-4A$^0$ distance were indicated.

#### Supplemental Figure S3.

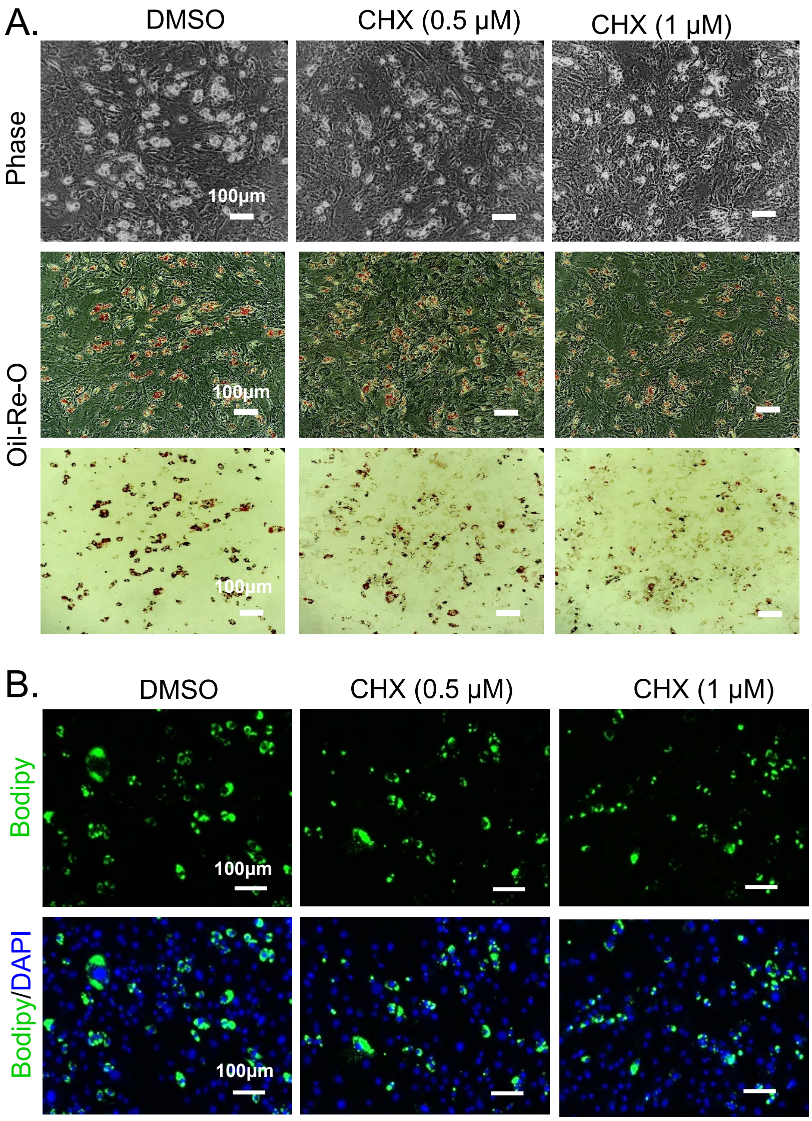


**Figure S3. Effect of Chlorhexidine on inhibiting early adipogenesis of primary preadipocytes.** (A) Representative images of phase-contrast and oil-red-O staining, and (B) Bodipy fluorescence staining of Primary preadipocyte at day 4 of early differentiation at indicated chlorhexidine (CHX) concentration. Scale bar: 100 μm.

#### Supplemental Figure S4.

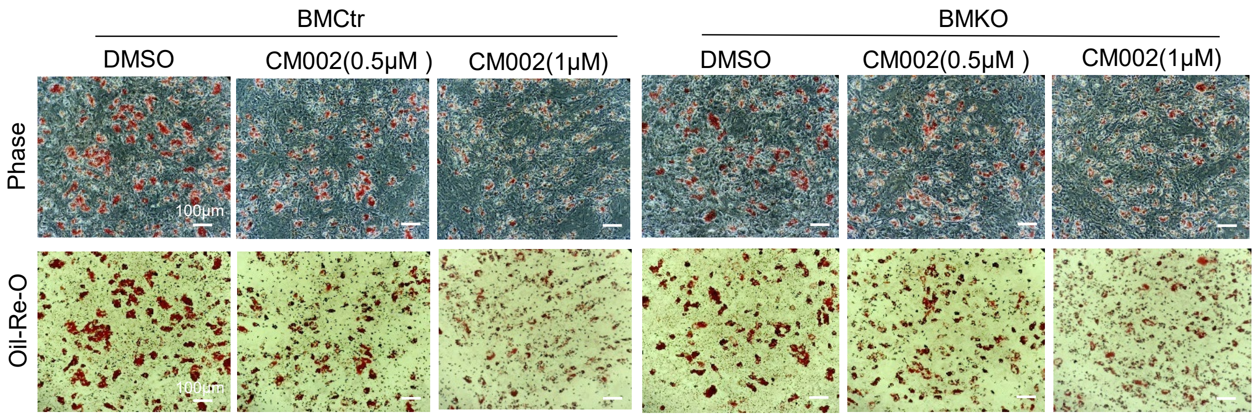


**Figure S4. Effect of CM002 on inhibiting terminal differentiation of primary preadipocytes from control (FloxCtr) or Bmal1-null (BMKO) mice.** Representative images of phase-contrast and oil-red-O staining with CM002 treatment at indicated concentrations at day 6 differentiation are shown. Scale bar: 100 μm.

### Supplemental Figure S5.

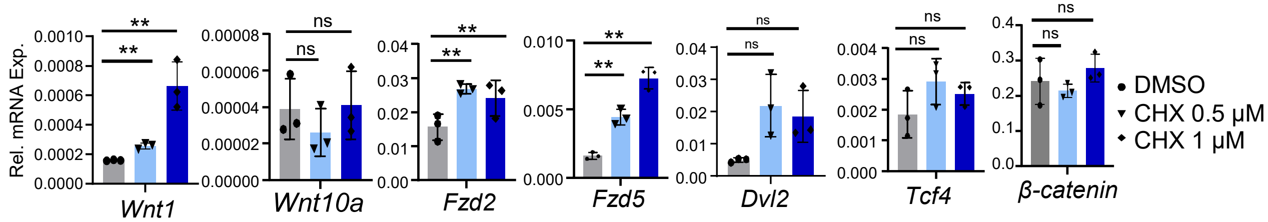


**Figure S5. Effect of CHX on inducing key components involved in Wnt signaling pathway in C3H10T1/2 mesenchymal precursor cells.** RT-qPCR analysis of mRNA expression of key components of Wnt signaling pathway of cells treated with indicated concentrations of chlorhexidine for 6 hours. N=3 replicates. *, **: p<0.05 and 0.01 CHX vs. DMSO by Student's t test.

#### Supplemental Figure S6.

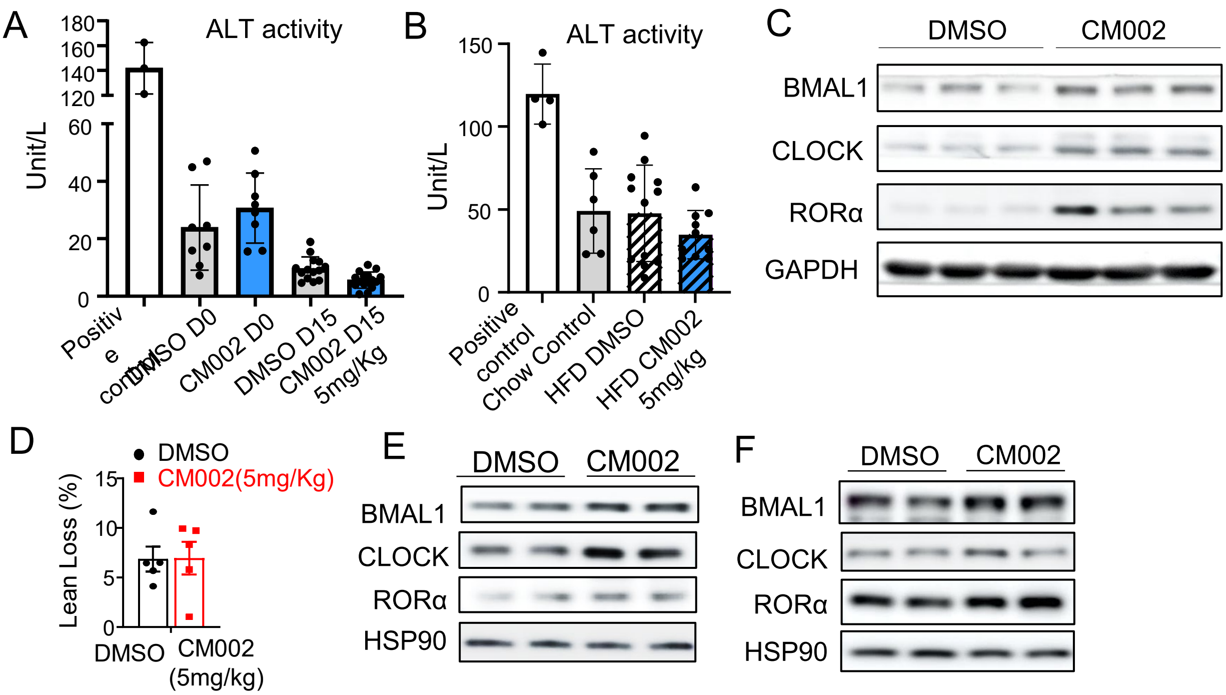


**Figure S6. Effect of in vivo intraperitoneal CM002 delivery in mice under chow or high-fat diet feeding condition.** (A, B) Plasma alanine aminotransferase (ALT) activity analysis of mice administered with 15 days of CM002 on chow diet (A) or for 7 days on high-fat diet (B). (C) Immunoblot analysis of clock protein expression in CM002-treated mice gastrocnemius muscle samples on regular chow diet. Each lane represent pooled sample of n=2 mice. (D, E) Immunoblot analysis of clock protein expression of inguinal WAT (iWAT, D) and brown adipose tissue (BAT, E) in CM002-treated mice on high-fat diet. Each lane represent pooled sample of n=3 mice. (F) The percentage of change in total lean mass by NMR analysis after 7 days of daily CM002 (5mg/Kg) treatment in mice on high-fat diet (n=5/group).

#### Supplemental Table 1. Primary antibodies list.

| Antibody | Source | Cat# | Dilution |
|---|---|---|---|
| C/EBPα | Santa Cruz | SC-61 | 1:1000 |
| C/EBPβ | Santa Cruz | SC-7962 | 1:1000 |
| PPARγ | Santa Cruz | SC-7273 | 1:1000 |
| FASN | Santa Cruz | SC-48357 | 1:1000 |
| FABP4 | Santa Cruz | SC-271529 | 1:1000 |
| PGC1α | Sigma | ST1204 | 1:1000 |
| BMAL1 | Santa Cruz | SC-365645 | 1:1000 |
| RORα | Proteintech | 82930-1-RR | 1:1000 |
| CLOCK | Cell Signaling | 5157S | 1:1000 |
| DBP | Proteintech | 12662-1-AP | 1:1000 |
| SREBP1 | Santa Cruz | SC-13551 | 1:1000 |
| GLUT4 | Cell Signaling | 2133S | 1:1000 |
| PDK4 | Proteintech | 12949-1-AP | 1:1000 |
| TOM20 | Santa Cruz | SC-17764 | 1:1000 |
| SDHB | Proteintech | 10620-1-AP | 1:1000 |
| UCP1 | Cell Signaling | 72298S | 1:1000 |
| β-Catenin | Cell Signaling | 8480S | 1:1000 |
| β-Actin | Proteintech | 66009-1-Ig | 1:3000 |
| HSP90 | Cell Signaling | 4874S | 1:3000 |

#### Supplemental Table 2. Primer sequence for qPCR analysis.

| Genes | | Sequences |
|---|---|---|
| Bmal1 | Forward | CGCTTTCTGGAGGGTGTCCGC |
| | Reverse | TGCCAGGACGCGCTTGTACC |
| Clock | Forward | TTGCTCCACGGGAATCCTT |
| | Reverse | GGAGGGAAAGTGCTCTGTTGTAG |
| Nr1d1 | Forward | TGGCATCCGGTGCACTGCAG |
| | Reverse | CCCTCCAGAAGGGTAGCACGCT |
| Nr1d2 | Forward | GGAGTTCATGCTTGTGAAGGCTGT |
| | Reverse | CAGACACTTCTTAAAGCGGCACTG |
| Cry1 | Forward | CTGGCGTGGAAGTCATCGT |
| | Reverse | CTGTCCGCCATTGAGTTCTATG |
| Cry2 | Forward | TGTCCCTTCCTGTGTGGAAGA |
| | Reverse | GCTCCCAGCTTGGCTTGA |
| Per1 | Forward | CTGCCATGGAGGAAGAAGAG |
| | Reverse | AGCTGGGGCAGTTTCCTATT |
| Per2 | Forward | ATGCTCGCCATCCACAAGA |
| | Reverse | GCGGAATCGAATGGGAGAAT |
| Dbp | Forward | CCACCGCGCAGGCTTGACAT |
| | Reverse | ACAGGGCGAGATCAGCGGGA |
| Wnt1 | Forward | GGTTTCTACTACGTTGCTACTGG |
| | Reverse | GGAATCCGTCAACAGGTTCGT |
| Wnt10a | Forward | CAACGCGTGCGCTCTGGGTA |
| | Reverse | TGGCTCAAGCCCTTTCCGCG |
| Fzd2 | Forward | CATGCCCAACCTTCTTGGC |
| | Reverse | CAGCGGGTAGAACTGATGCAC |
| Fzd5 | Forward | AATCATGCAGGGGGCCCCGAA |
| | Reverse | CGACAAGCTAGGTACCTGTGGCG |
| Dvl2 | Forward | TCAGTTTGCGGGTGTGCGCAG |
| | Reverse | TTCGTCTCGCCTACACCACCG |
| Tcf4 | Forward | GGCCGCAGCGCCTTCTCTTTA |
| | Reverse | ACCATCATTGACTCCCCCGAGG |
| Ucp1 | Forward | TAACGGGTCCTCCCTGCCCG |
| | Reverse | CCGCGACTTCGGACTCCTGC |
| Dio2 | Forward | GATGGCTGGGCAGTGCCTGG |
| | Reverse | GGGCGGCAAGGAGAAACGCT |
| Hsl | Forward | TGTTGGGGTGACTCTAACGC |
| | Reverse | GAGCTCCGCCTTTAATGGGT |
| β-catenin | Forward | CGCTTGGCTGAACCATCAC |
| | Reverse | GTTCCGCGTCATCCTGATAGT |
| 36B4 | Forward | CGCTTTCTGGAGGGTGTCCGC |
| | Reverse | TGCCAGGACGCGCTTGTACC |
